## Supplementary material for "Pooled genome-wide CRISPR activation screening for rapamycin resistance genes in *Drosophila* cells": Dataset S1

Supplementary file 1. sgRNA vectors used in this study.

| vector | description | 1st sgRNA sequence | 2nd sgRNA sequence |
|---|---|---|---|
| sg*Jon25Biii* | Single sgRNA vector | GAAATCAAAGAAACTCTAGG | |
| sg*Sdr* | Single sgRNA vector | AGAATGAAAATCAAAGGTGG | |
| sg*CG9877* | Dual-sgRNA vector | ATATTGTGTGGATATTGTAT | CTAATACGAACGCGATATTT |
| sg*CG8468* vector 1 | Dual-sgRNA vector | ACACTGCTAGAACGTGTTGT | TCCGTCGCTACTCCATAGTG |
| sg*CG8468* vector 2 | Dual-sgRNA vector | CGTATACGACACAGTGTTCT | ACACTGCTAGAACGTGTTGT |
| sg*CG8468* vector 3 | Dual-sgRNA vector | GATGAAGGTGCTATTTGGTA | CTTCGGATTTGTTGGTTACA |
| sg*CG8468* vector 4 | Dual-sgRNA vector | CTTCGGATTTGTTGGTTACA | CGTATACGACACAGTGTTCT |
| sg*CG8468* vector 5 | Dual-sgRNA vector | TCCGTCGCTACTCCATAGTG | TGTTGGCAGTATAAACCGCG |
| sg*CG8468* vector 6 | Dual-sgRNA vector | TGTTGGCAGTATAAACCGCG | GATGAAGGTGCTATTTGGTA |
| sg*CG5399* vector 1 | Dual-sgRNA vector | TTGCAGGTAAGACAACCTAT | AAAAGATACAAGCCTATAAA |
| sg*CG5399* vector 2 | Dual-sgRNA vector | TGGAATGACTAATCCCCACG | TGGACACCGAATCTATTTCT |
| sg*CG5399* vector 3 | Dual-sgRNA vector | GTTTGTTTATACCTTTCAAG | TGGAATGACTAATCCCCACG |
| sg*CG5399* vector 4 | Dual-sgRNA vector | TGGACACCGAATCTATTTCT | GCCAGGCGCTCCACAAGTGC |
| sg*CG5399* vector 5 | Dual-sgRNA vector | AAAAGATACAAGCCTATAAA | GTTTGTTTATACCTTTCAAG |
| sg*CG5399* vector 6 | Dual-sgRNA vector | GCCAGGCGCTCCACAAGTGC | TTGCAGGTAAGACAACCTAT |
| sg*CG9932* vector 1 | Dual-sgRNA vector | GCCGAAGTCGCACGAACAAC | GAGCTGTTGGGTGTGCGGTT |
| sg*CG9932* vector 2 | Dual-sgRNA vector | GAGCTGTTGGGTGTGCGGTT | CTTCTTCGCTCTCTTAAGCG |

| | | | |
|---|---|---|---|
| sg*CG9932* vector 3 | Dual-sgRNA vector | AACGGCCGGTGCTAACAGAG | GCCGAAGTCGCACGAACAAC |
| sg*CG9932* vector 4 | Dual-sgRNA vector | TGCCACCGACTGGTTGGCGT | AACGGCCGGTGCTAACAGAG |
| sg*CG9932* vector 5 | Dual-sgRNA vector | CTTCTTCGCTCTCTTAAGCG | AGCGCGAGAGCAAGCGAACG |
| sg*CG9932* vector 6 | Dual-sgRNA vector | AGCGCGAGAGCAAGCGAACG | TGCCACCGACTGGTTGGCGT |

Supplementary file 2. PCR primers used in this study.

| primer | description | Sequence (5'-3') |
|---|---|---|
| Focused-NGS-1F | 1st round PCR of NGS library for focused screen | CCTATTTTCAATTTAACGTCG |
| Focused-NGS-1R | 1st round PCR of NGS library for focused screen | CTAGCCTTATTTTAACTTGC |
| Focused-NGS-2F | 2nd round PCR of NGS library for focused screen | CCTACACGACGCTCTTCCGATCT-$(N)_n$-$(B)_6$-CCTATTTTCAATTTAACGTCG |
| Focused-NGS-2R | 2nd round PCR of NGS library for focused screen | CTAGCCTTATTTTAACTTGC |
| Focused-NGS-3F | 3rd round PCR of NGS library for focused screen | AATGATACGGCGACCACCGAGATCTACACTCTTTCCCTACACGACGCTCTTCCGATCT |
| Focused-NGS-3R | 3rd round PCR of NGS library for focused screen | CAAGCAGAAGACGGCATACGAGATCTAGCCTTATTTTAACTTGC |
| WGS-NGS-1F | 1st round PCR of NGS library for genome-wide screen | CCTATTTTCAATTTAACGTCG |
| WGS-NGS-1R | 1st round PCR of NGS library for genome-wide screen | ATATGCTTTATTGACAGAAAATTTGATG |
| WGS-NGS-2F | 2nd round PCR of NGS library for genome-wide screen | CCTACACGACGCTCTTCCGATCT-$(N)_n$-$(B)_6$-CCTATTTTCAATTTAACGTCG |
| WGS-NGS-2R | 2nd round PCR of NGS library for genome-wide screen | ATATGCTTTATTGACAGAAAATTTGATG |
| WGS-NGS-3F | 3rd round PCR of NGS library for genome-wide screen | AATGATACGGCGACCACCGAGATCTACACTCTTTCCCTACACGACGCTCTTCCGATCT |
| WGS-NGS-3R | 3rd round PCR of NGS library for genome-wide screen | CAAGCAGAAGACGGCATACGAGATATATGCTTTATTGACAGAAAATTTGATG |
| q*Jon25Biii*-F | qPCR primer for *Jon25Biii* | CAAGCTGGTGGGAGTTAGCA |
| q*Jon25Biii*-R | qPCR primer for *Jon25Biii* | GGTCACGGATCCAGTCCAAG |
| q*Sdr*-F | qPCR primer for *Sdr* | CGGCTACTTCCAGACGCTAC |
| q*Sdr*-F | qPCR primer for *Sdr* | TGGCCACCAGTGAAGAAGA |
| q*CG9877*-F | qPCR primer for *CG9877* | GCGAGCTGCTATCGTGTTTG |
| q*CG9877*-R | qPCR primer for *CG9877* | CCTCCGAAGCCACCATATCC |
| q*CG13538*-F | qPCR primer for *CG13538* | GTTGGCCGATGAATTTGAAACG |
| q*CG13538*-R | qPCR primer for *CG13538* | ATCAGCCGATTGCGATACTTC |
| q*CG8468*-F | qPCR primer for *CG8468* | CTGATGGCAGAGTTCGGTGTG |

| | | |
|---|---|---|
| q*CG8468*-R | qPCR primer for *CG8468* | CATGGCACTAACAAAGGGTCC |
| q*CG5399*-F | qPCR primer for *CG5399* | AGTGCGATCATCTGCCTGG |
| q*CG5399*-R | qPCR primer for *CG5399* | TTGTGGCGTTGTTTGGGCT |
| q*CG9932*-F | qPCR primer for *CG9932* | TTGCCGTTGTGAACAGCGT |
| q*CG9932*-R | qPCR primer for *CG9932* | CGCTTGATCTTGCATTGCGG |

Supplementary file 3. dsRNAs used in this study.

| dsRNA | dsRNA sequence |
|---|---|
| ds*GFP* | AAGTTCATCTGCACCACCGGCAAGCTGCCCGTGCCCTGGCCCACCCTCGTGAC<br>CACCCTGACCTACGGCGTGCAGTGCTTCAGCCGCTACCCCGACCACATGAAGC<br>AGCACGACTTCTTCAAGTCCGCCATGCCCGAAGGCTACGTCCAGGAGCGCACC<br>ATCTTCTTCAAGGGCGACGGCAACTACAAGACCCGCGCCGAGGTGAAGTTCGA<br>GGGCGACACCCTGGTGAACCGCATCGAGCTGAAGGGCATCGACTTCAAGGAGG<br>ACGGCAACATCCTGGGGCACAAGCTGGAGTACAACTACAACAGCCACAACGTCT<br>ATATCATGGCCGACAAGCAGAAGAACGGCATCAAGGTGAACTTCAAGATCCGCC<br>ACAACATCGAGGACGGCAGCGTGCAGCTCGCCGACCACTACCAGCAGAACACC<br>CCCATCGGCGACGGCCCCGTGCTGCTGCCCGACAACCACTACCTGAGCACCCA<br>GTCCGCCCTGAGCAAAGA |
| ds*CG5399*-1 | ATGTGCAATAGTGCGATCATCTGCCTGGCAATCGTGGCCGCCGTGGCCTCCATC<br>TCTCAGGTTTCGGCCCAAGCCCAAACAACGCCACAATCCTGCGCCCAAGCCACC<br>GGCATTAGTTTTAACCAGACCGCT |
| ds*CG5399*-2 | TTTAGTGGTCAGTGGGTCGAGTTGGCCCGCAATCCCGCCTCAAATGTCTCCTGC<br>ATTTCCCTTAACGTGGCTTTCTGGAACAACAACAATTACATGCTGGTCAATGTCA<br>GCCATTCAAGCAATGTCGCCTCCACTTTGGTGGACGTCTATGAAACTGCGAATAT<br>CACGCTGGTGGAGAACAACCTCAGCGGATACAATGTCACCCTGGTGAATGGCCA<br>GAAGCAGGAGTACGTCTTCGTGAAGTTGCTTAGCTTTGTGAACTCCACCTATCTG<br>GTTGGATGCACCTACACCAATGCTACCAACACATCAACCAGTGCTGGTTTTATTC<br>TGGGTAGGTCCACCTATACTCCGGAAGGCGTAAGGATTGCCAACAATGATGCCT<br>CCGTGTCCTTCAGCAATTTCCAGAACAACACCTACGGCAATGTCACTCAGTTGGA |
| ds*Pi3K92E*-1 | CCTACCGCGGATGACACGCCTCTGCATTGTGATCTTCGAGGTGACCAAGATGTC<br>GCGCTCGAAGAAATCGTCCAACAATAAGGACATCGCTCTGAAAGACGTCCCGTA<br>CAACAAAAACCCGCTGGCCTGGGTCAACACAACAATTTTCGACCACAAGGACAT<br>CCTGCGCACCGGTCGCCATACGCTGTACACGTGGACGTATGCTGACGACATTCA<br>GTCGGTTGAGGTTTTCCATCCGCTGGGCACCATCGAACCCAATCCGCGCAAAGA<br>GGAATGTGCGCTGGTGGACCTCACATTCCTCAGCAGCGGAACCGGAACGGTGC<br>GGTATCCCAGCGAGGAGGTCGTGTTGCAGTACGCTGCGGATCGAGAGCAGGTC<br>AATCGACTGCAACGGCAACTGGCAGGGCCAGAAAAGCCGATCAAAGAACTGAA<br>GGAGCTTATGGCCAATTATACAGGACTGGATAAGATCTATGAGATGGTTGACCA<br>GGACCGCAATGCCATT |
| ds*Pi3K92E*-2 | AGAGATTGGCGTGCTGATTTAAATGGAAAACTTTGAAAACAACAATCTGTTGATA<br>AGGTAACCAGCGAAATGCGGGACAACTAGCCTCCGTAGCCTGAAAGAGGCCATT<br>GAAAAGTCTAATTAGGAGGAGCTGGTGATCCTCAAGATCCGAGGCACCAGATCC<br>AAAATCCGAACTTGAAGATCAAATAGTCATGAACATGATGGACAACCGGGCGTT<br>GGCCTACGTGGCCCACCAGCCCAAGTATGAGACACCGCCGGAAGAAGCGGAGC<br>CGCCCTGCATGCGCTTTTCGGTTAACCTGTGGAAAAACGAGATGCTGAACTGGG<br>TGGACCTAATCTGCCTGTTGCCCAATGGATTCCTGCTGGAGCTCAGGGTCAATC<br>CGGCCAACACCATCCAGGTAATCAAGGTGGAGATGGTCAACCAGGCCAAACAG<br>ATGCCACTGGGCTATGTGATCAAAGAGGCCTGCGAGTACCAGGTGTACGGCATC<br>TCGACCTTCAACATCGAACCTTACACCGACGAAACGAAGCGACTCAGTGAGGTC |
| ds*InR*-1 | GCCATGGAAAATGACCTGCCAGCCACAACGCCTACCAAGAAAATATCAGATCCT<br>TTAGCAGGCGACTGTAAGTGCGTGGAGGGTTCGAAGAAGACTAGCAGTCAGGA<br>ATACGATGATCGTAAAGTTCAAGCGGGCATGGAGTTTGAGAACGCGTTGCAAAA<br>CTTTATATTTGTTCCAAACATTCGGAAAAGCAAGAATGGATCGTCTGACAAATCA<br>GACGGAGCGGAAGGTGCAGCTCTCGATTCTAATGCTATTCCAAATGGAGGAGCT<br>ACTAACCCTTCACGTAGAAGGAGAGACGTTGCGCTCGAGCCAGAGCTCGACGAT<br>GTAGAGGGCAGTGTACTTCTACGCCATGTGCGCTCCATCACAGACGATACCGAT<br>GCATTTTTCGAAAAGGACGACGAAAATACCTATAAAGACGAAGAAGACTTGTCCT<br>CCAACAAACAATTCTATGAGGTGTTTGCCAAGGAATTGCCACCAAATCAAACACA<br>TTTTGTCTTTGAAAAACTGCGCCACTTCAC |
| ds*InR*-2 | GGAACAGCCGTTCTCAATGTCACATTACAATCAGTGGGAGCAAACTCCGCTATG<br>CTGAACGTCACGACAAAAGTTGAAATAGGAGAGCCCCAAAAGCCGAGCAATGCT |

| | |
|---|---|
| | ACAATTGTTTTTAAGGATCCGCGCGCCTTCATCGGTTTCGTGTTTTATCATATGATCGATCCGTACGGGAACTCAACTAAAAGCAGTGACGATCCATGCGATGATCGCTGGAAGGTTAGCTCTCCGGAAAAGAGCGGGGTCATGGTATTAAGCAATTTGATTCCGTACACTAACTACTCCTACTACGTTCGGACCATGGCTATATCCTCGGAATTGACAAACGCGGAGAGCGACGTGAAGAACTTTAGGACGAATCCCGGACGACCGTCAAAGGTTACGGAGGTGGTAGCAA |
| ds*Pvr*-1 | TCGGGTCATTACAACGTTCAGGAATATGCCAATCGCACGATCCAAATGACCGCGAACTTTGAGGGATTTCCGACGCCCTCCTTCAGTTGGTTCAAACCCGATGGCACCGAGGTGCGACAATCGGAGAATAACTTCAAGATTCTCTCCACGGAATTGAGCACAATGCTCCAGGTGCTGAACGCCCAATTGCAGGACAGCGGCACGTATGTCCTCCGTGGATCCAATTCCTTCGGCGTCGTTCAGCGGGAGTACAACGTCAGTGTGATGGACGCACCGGCGCTGAAGATGTCGGACGCCTATGTCCAGGTGGGATCCGTGGCGCGACTGGAGTGCACAGTACGCTCCTATCCGCCGGCTATCGTGACCTTCTT |
| ds*Pvr*-2 | CAGCAGCCGGGAGATAGACTTGTACGTCCACGATCCCTCTGCTCCTCAGTGGACAAACGGCGGACAGGAGGGTCACTCGAAAATAAAGCGCAAACTAAGCCAAACGCTGGAGCTGGAGTGTGCCTCCACAGCGGTTCCCGTGGCAATTGTGCGTTGGTTTAAGGACGACAAGGAAGTGACCGAATCAAAGCTCAGGCACATCATTGAAAAGGAATCCAAGCTGCTGATCACTCACCTGTATCCCGGAGATGAAGGCGTCTACAAGTGTGTGGTGGAGAACCGATTGGACAGAATCGAACGCTCCTTCACGGTAGTGATATCAGATCTGCCCGGCATTAGCATGGCCTGGGTGTGGTTCGGTGTGATACTATTCCTCATCCTGATCGGTCTGTGCGTCTTCCTCGCCGTGCGCTACCAGAAGGAGCACAAGCGGCATCTGGCCCTTAAGGCAGCCGGATTGGCCAACTTCGAGGAGGGCGCCGTGGGACACATCAATCCCGATCTGAC |
| ds*Flo1*-1 | TACCTGAGGTCATTGGGTATGGCCCGCACGGCGGAGGTGAAGCGCGATGCCCGCATTGGCGAGGCTGAGGCCCGAGCGGAGGCCCACATTAAGGAGGCCATTGCCGAGGAGCAACGCATGGCCGCACGCTTCCTCAACGATACCGATATTGCCAAGGCCCAGCGCGACTTTGAGCTGAAGAAGGCAGCATACGATGTGGAAGTGCAGACCAAAAAGGCCGAGGCCGAGATGGCCTACGAGCTGCAAGCGGCCAAGACCAAGCAGCGCATCAAGGAGGAGCAGATGCAGGTGAAGGTGATCGAGCGCACGCAGGAGATTGCCGTCCAGGAGCAGGAGATCATGCGCCGCGAGCGAGAGCTGGAGGCCACCATCCGCCGACCGGCCGAGGCGGAGAAGTTCCGCATGGAGAAATTGGCCGAGGCCAACAAGCAGCGCGTGGTCATGGAAGCCGAG |
| ds*Flo1*-2 | CCAGGTAGAGAGTCCTTGCGTGTACACCAGCCAAGGAGTGCCCATCTCGGTGACAGGCATTGCACAGGTGAAGGTCCAGGGTCAGAACGAGGACATGCTGCTGACCGCCTGTGAGCAGTTCTTGGGCAAATCAGAGGCAGAGATCAACCACATCGCCTTGGTCACCCTGGAGGGGCATCAGCGTGCCATCATGGGTTCGATGACCGTGGAGGAGATCTACAAGGACCGCAAGAAGTTCAGCAAGCAGGTGTTTGAGGTGGCCTCCAGCGATTTGGCCAACATGGGAATAACCGTGGTTTCCTACACCATC |
| ds*Flo2*-1 | TGCTGTGGATCGACCAAGAAGCGCACGATTGTGGGCGGCTGGGCGTGGGCGTGGTGGCTGGTAACCGATGTCCAGCGACTGTCCCTCAATGTGATGACCCTGAATCCGATGTGCGAGAATGTGGAAACGTCGCAAGGTGTTCCGCTAACGGTGACCGGAGTGGCTCAATGCAAGAT |
| ds*Flo2*-2 | GCGAGGTGGCCGCACCGGACGTGGGTCGTATGGGCATCGAGATTCTCTCGTTTACGATCAAGGACGTCTACGATGATGTGCAGTACCTGGCCTCGTTGGGCAAGGCCCAGACCGCCGTGGTCAAGCGGGATGCAGATGCCGGCGTGGCGGAGGCCAATCGAGATGCCGGTATCCGTGAGGCGGAGTGCGAAAAGAGCGCCATGGATGTGAAATACTCGACGGACACGAAAATCGAGGACAACACCAGGATGTACAAGCTGCAGAAGGCCAATTTCGATCAGGAGATCAACACGGCCAAGGCCGAATCGCAGTTGGCCTACGAGCTGCAGGCAGCCAAGATCCGCCAGCGCATCCGTAACGAGGAGATTCAGATCGAGGTGGTGGAGCGACGCAAGCAGATCGAGATTGAGTCGCAGGAAGTGCAGCGCAAGGATCGCGAGCTCACTGGCACAGTCAAGCTGCCCGCCGAGGCCGAGGCCTTCCGCCTCCAGACCCTTGCGCAGGCCA |
| ds*Chc*-1 | TGTGCGAAAGTTCAACAAGCTCTTTACAGCCGGCCAGTATGCTGAAGCGGCTAAAGTTGCTGCCCTGGCACCCAAGGCCATTCTGCGTACGCCACAGACGATCCAGCGTTTCCAACAGGTGCAGACACCAGCTGGCTCCACGACTCCGCCGCTGCTGCAAT |

| | |
|---|---|
| | ACTTTGGCATTCTCCTCGACCAGGGCAAGCTGAACAAGTTCGAGTCTCTCGAGCTGTGCCGTCCCGTCTTGCTGCAGGGCAAGAAGCAGCTGTGCGAGAAGTGGCTGAAGGAGGAGAAGTTGGAATGCAGCGAGGAGTTGGGTGATCTGGTCAAGGCCTCCGATCTTACACTTGCCCTGTCCATCTATCTGCGCGCAAATGTGCCCAACAAGGTTATCCAATGCTTTGCTGAGACTGGGCAGTTCCAGAAGATTGTACTCTACGCCAAGAAGGTCAACTATACGCCCGATTACGTGTTCCTGCTGCGCTCCGTGATGCGAAGCAACCCGGAGCAAGGAGCTGGTTTCGCCTCTATG |
| ds*Chc*-2 | CCCGAACGGGTGAAGAACTTCTTGAAGGAGGCCAAGCTGACGGATCAGCTACCATTAATTATTGTTTGTGATCGTTTTGATTTCGTGCACGACTTGGTGCTTTACCTGTATCGTAACAATCTGCAGAAGTACATTGAGATCTATGTGCAGAAAGTGAATCCATCCCGCTTGCCAGTGGTAGTGGGTGGTCTTCTTGATGTTGATTGCAGTGAGGATATAATTAAAAATCTAATTCTCGTGGTCAAGGGACAATTCTCAACCGACGAACTGGTCGAGGAGGTCGAGAAGCGCAACCGTCTCAAGCTTCTCCTTCCCTGGCTGGAGTCCCGAGTTCACGAGGGCTGCGTCGAGCCAGCCACCCACAACGCGTTGGCCAAGATCTACATTGACTCGAACAACAATCCCGAGAGATATCTTAAGGAGAATCAGTACTACGATAGCCGTGTGGTCGGTCGCTACTGCGAGAAGCGGGATCCCCATTTGGCGTGTGTCGCCTACGAGCGTGGATTGTG |
